# Widespread but cryptic introgression shapes genetic diversity in natural populations

**DOI:** 10.64898/2026.07.06.736689

**Authors:** Guillaume Lavanchy, Ludovic Ruedi, Olivier Broennimann, Kristine Jecha, Marianna Tzivanopoulou, Jérôme Goudet, Tanja Schwander

**Affiliations:** Department of Ecology and Evolution, University of Lausanne, CH-1015 Lausanne, Switzerland; Biology Department, Lund University. Ekologihuset, Sölvegatan 37, 223 62 Lund, Sweden

**Keywords:** *Camponotus*, *Formica*, *Lasius*, *Myrmica*, *Tapinoma*, *Temnothorax*, *Tetramorium*, hybridization

## Abstract

Introgression following hybridization is increasingly recognized as a major driver of evolution. However, its importance depends on its frequency in nature, which remains to be quantified. To address this, we provide a snapshot of ongoing introgression in a whole species assemblage (4126 ant colonies). 23% of all 82 local species show signs of introgression, which is more than twice previous estimates. Introgression is typically subtle, yet contributes measurably to genetic diversity. Species divergence, rather than classical prezygotic reproductive barriers (mating phenology, ecological niche, fine-scale habitat use) constrains introgression, suggesting that the main reproductive barriers are postzygotic at this stage of divergence. Our results indicate that introgression may be a common but often overlooked feature of natural communities.

## Introduction

Introgression, the exchange of genetic material between species following hybridization, is increasingly recognized as a major driver of evolution (*1, 2*). It can reshape species boundaries (*3–5*) contribute to adaptation (*6*), and, in some cases, generate new species (hybrid speciation; *7*). Despite its well-documented evolutionary consequences, a central question remains: how common is introgression in natural populations? Indeed, the evolutionary impact of introgression depends not only on the nature and magnitude of its effects, but also on how frequently it occurs. Yet, we currently lack unbiased estimates of how frequent introgression is within natural communities.

Most existing estimates rely on the proportion of hybridizing species and serendipitous reports of hybrids. Together, they suggest that about 9% of plants and 10% of animal species hybridize (*8, 9*), but these figures vary widely across taxa and are inherently biased toward conspicuous or well-studied systems (e.g., *10–13*). Moreover, such estimates focus on whether species hybridize at all, rather than how often hybridization occurs and leads to introgression, which is what ultimately determines its evolutionary relevance.

Attempts to quantify hybridization rates within species are similarly limited. They typically focus on known hybridizing taxa or on hybrid zones, leading to substantial overestimates. Morphology-based surveys (e.g., *14–17*) likely underestimate hybridization because hybrids are difficult to detect and rapidly become indistinguishable from parental forms after backcrossing. Thus, it remains unknown how pervasive hybridization is in natural populations.

Recent genomic approaches have revealed widespread signatures of ancient introgression across many taxa (*18–21*). However, these retrospective signals reflect not only the occurrence of hybridization but also its subsequent filtering by selection and drift (*2*), and therefore provide limited insight into the frequency and drivers of ongoing gene flow. What is needed instead is a population genomic approach that directly quantifies contemporary introgression across entire communities. By genotyping large numbers of individuals from multiple co-occurring species, such an approach can provide an unbiased snapshot of introgression at a given time and place.

In this study, we use such a population genomic approach to quantify contemporary introgression across an entire species assemblage. We leverage a high-resolution survey of co-occurring ant species within a natural or semi-natural landscape to obtain an unbiased estimate of the prevalence of ongoing gene flow across species, based on 4126 individuals from 40 species. We then assess how much introgression contributes to standing genetic variation within species and test which barriers (prezygotic or postzygotic) most strongly limit gene flow. This framework allows us to move from anecdotal and retrospective evidence toward a direct, community-level view of introgression in nature.

## Results and Discussion

We surveyed ant diversity across the canton de Vaud, Switzerland, sampling 8,841 colonies and identifying 82 species from 21 genera based on morphology. To quantify introgression at the community level, we focused on the subset of genera represented by at least two species and 10 sampled colonies per species (*Camponotus, Formica, Lasius, Myrmica, Tapinoma, Temnothorax, Tetramorium*). From these, we genotyped one worker per colony, yielding 5,144 individuals. Genotyping was done using ddRAD sequencing and complemented with barcoding of the COI mitochondrial gene. After stringent filtering and removal of potential cross-contaminations, we corroborated morphological species identification using SNP-based admixture analyses, multidimensional scaling, and COI phylogenies (Figures S1 - S7). The final dataset post-filtering contained 4126 individuals from 40 species distributed across the study area (Figure 1A).

**Figure 1.**
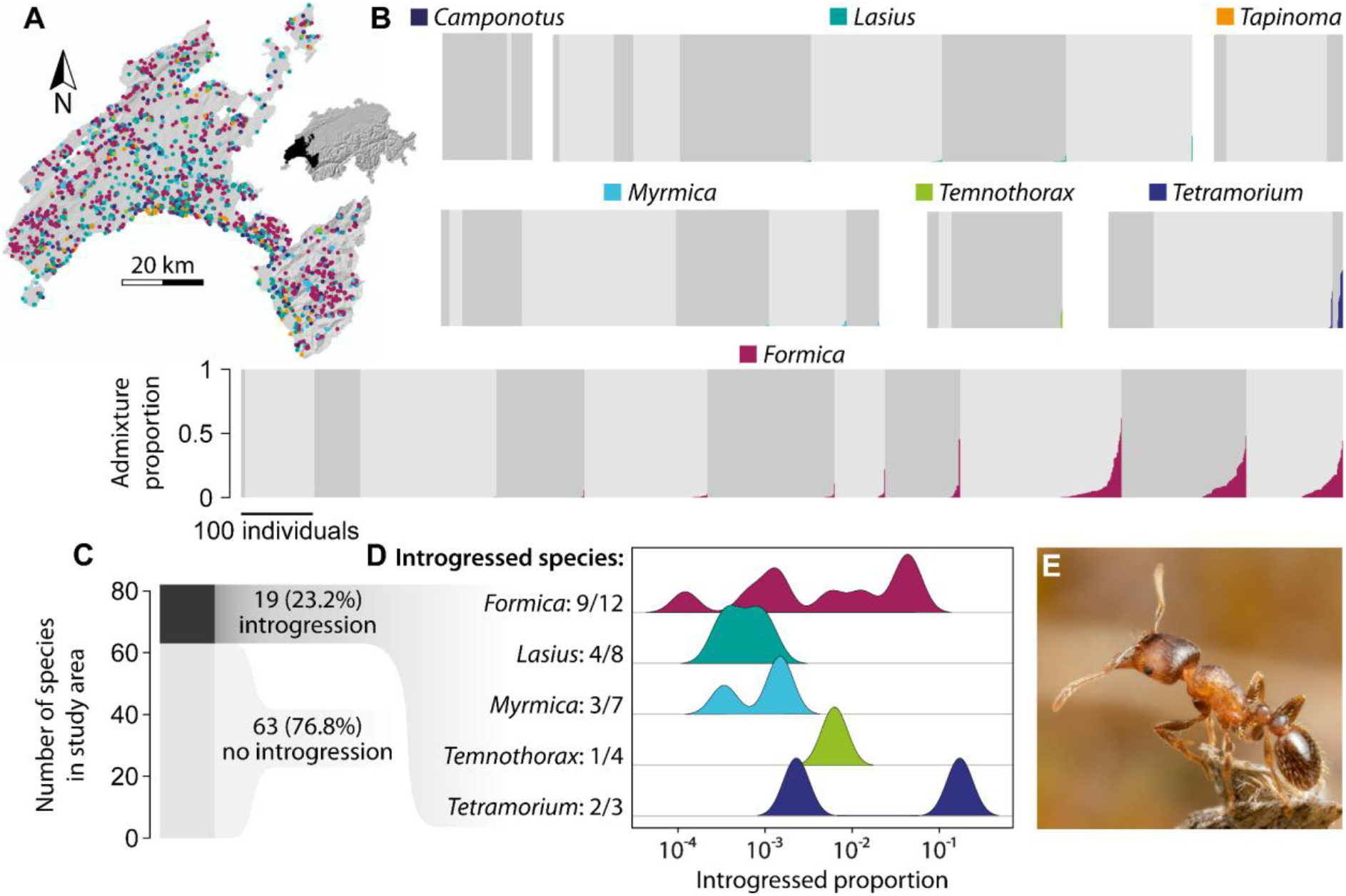
**A:** Map of the studied area with locations of the 4126 retained samples, colored by genus. Inset: Switzerland, with the study area (canton de Vaud) in black. **B:** Admixture proportion for each of the 4126 samples. Alternating shades of grey distinguish different species, non-admixed; colored shades indicate admixture from other species. Species within genus and individuals within species are sorted from least (left) to most (right) introgressed. A version with individual colors for each species is available as Figure S8. **C:** Barplot of introgressed or non-introgressed species sampled in the study area. Note that “no introgression” includes non-genotyped species (fewer than two species and 10 samples). **D:** Number of species per genus showing signatures of introgression and distribution of introgression proportion by species, measured as the total amount of genomic material of all individuals from one species derived from other species (i.e. the proportion of the colored area in **B**). Note the logarithmic scale of the x-axis. **E**: A worker of *Tetramorium sp*., a genus where high introgression was detected. Credit Bart Zijlstra, bartzijlstra.com.

We quantified ongoing admixture to obtain a snapshot of “contemporary introgression”, defined here as gene flow recent enough that introgressed alleles are still segregating within populations. The admixture proportions for all 4126 individuals are shown in Figure 1B. Strikingly, 23.2% of all species showed clear signatures of introgression (Figure 1C). This is more than double previous estimations of hybridization for animals (*9*) and plants (*8*). Note that the proportion of hybridizing species could be substantially higher as only the fraction of hybridization events that lead to viable and fertile offspring and further backcrosses lead to introgression.

The relevance of introgression for species evolution not only depends on its frequency in nature, but also on its contribution to genetic variation (*6*). To assess this contribution, we estimated nucleotide diversity (π) within each species that engaged in hybridization, including and excluding admixed individuals. We found that introgression increased nucleotide diversity for most species (mean = 2.5%; range: -0.16% – 29%), with an exceptional case (*Tetramorium immigrans*, the species showing the highest introgressed proportion) where introgression increased nucleotide diversity by 29% (Figure 2). The significance of a 2.5% average increase in genetic diversity is not immediately intuitive, but a comparison is provided by the fact that genetic diversity scales with census population size across the tree of life (*22*). Based on this relationship, a 2.5% increase in genetic diversity corresponds to a much more substantial 21% increase in population size. In other words, genetic diversity contributed by introgression amounts to the difference in diversity between two species which would differ in population size by 21%.

**Figure 2.**
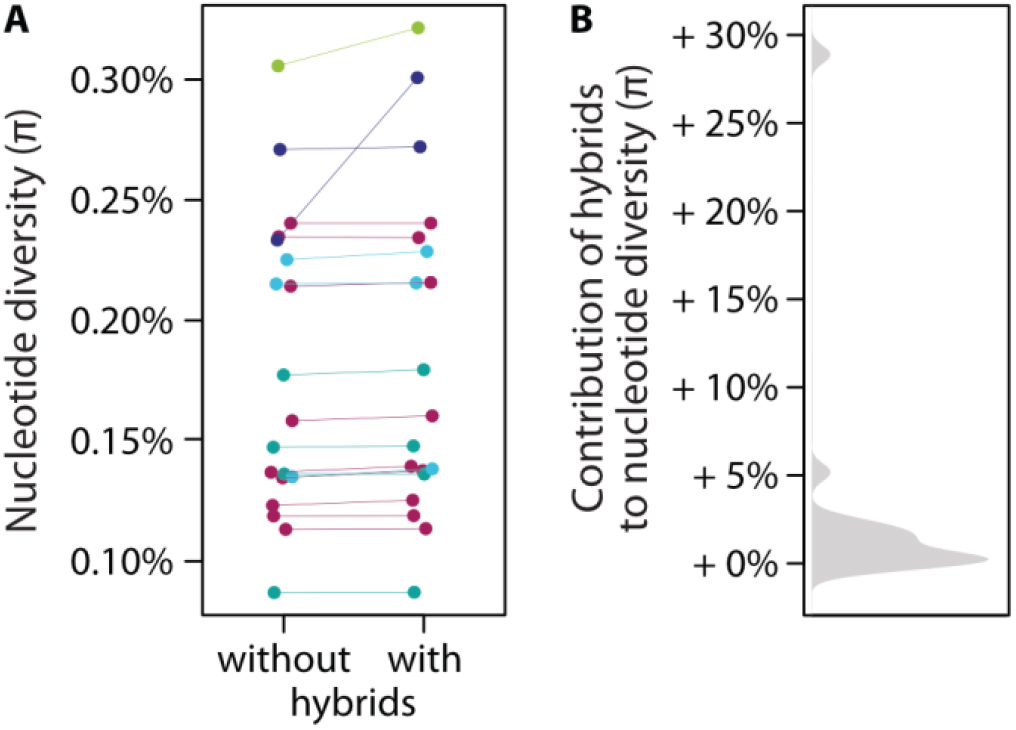
The contribution of introgression to nucleotide diversity. **A:** Each pair of points is one species with a signature of introgression. Nucleotide diversity excluding (left) and including (right) all individuals showing signs of introgression. Colors according to genus, as in **Figure 1. B:** Distribution of the relative contribution of introgressed individuals to nucleotide diversity, computed as the ratio of the right to left values in **A**.

What is the evolutionary relevance of this increased genetic diversity? Variants entering a population via introgression could contribute disproportionately to adaptation compared to *de novo* mutations. First, introgressed variants have already escaped purging in the donor species and are therefore less likely to be strongly deleterious. Second, they may enter the population as haplotypes consisting of several linked variants, potentially causing larger molecular and phenotypic changes, which can facilitate the crossing of adaptive valley (*6*). Consistent with this, introgression has been associated with rapid adaptation, in some cases allowing hybrid lineages to outpace non-admixed populations (*23–25*).

On the other hand, whether neutral genetic variation is a good measure of evolutionary potential is debated (*26*) and complementing genetic diversity with phenotypic studies will be needed to assess the net impact of introgression on adaptation. Many introgressed variants are likely to be involved in incompatibilities and thus to be deleterious. This is because introgressed variants enter a novel genomic background, which results in the mixing of coevolved regulatory networks and can lead to transgressive and potentially deleterious expression patterns (*27, 28*). In addition, they get exposed to the environment of the receiving species, to which they may not be locally adapted. However, highly deleterious variants (i.e. those involved in incompatibilities or hybrid load) are expected to be purged quickly from hybrid genomes. Since most of the hybrids we discovered were the result of several generations of backcrosses (Figure 1B), such highly deleterious variants should have been eliminated already, and their contribution to the increased diversity we detected is likely limited. Overall, the net effect of introgression on adaptation thus depends on locus-specific patterns of expression and regulation, and on the level of divergence between the parental species.

We found a strong negative relationship between genetic divergence (d_XY_, a proxy for incompatibilities) and introgression (Figure 3; p < 10^-5^), consistent with postzygotic barriers limiting gene flow. This raises the question of whether prezygotic barriers play a comparable role in the surveyed natural populations. To test this, we quantified three classical prezygotic barriers: differences in mating phenology, ecological niche divergence, and fine-scale spatial co-occurrence, and asked whether they predict introgression rates. We found that none of these were associated with introgression proportions (Figure 3, Figure S9; permutation LM: all *p* > 0.2).

**Figure 3.**
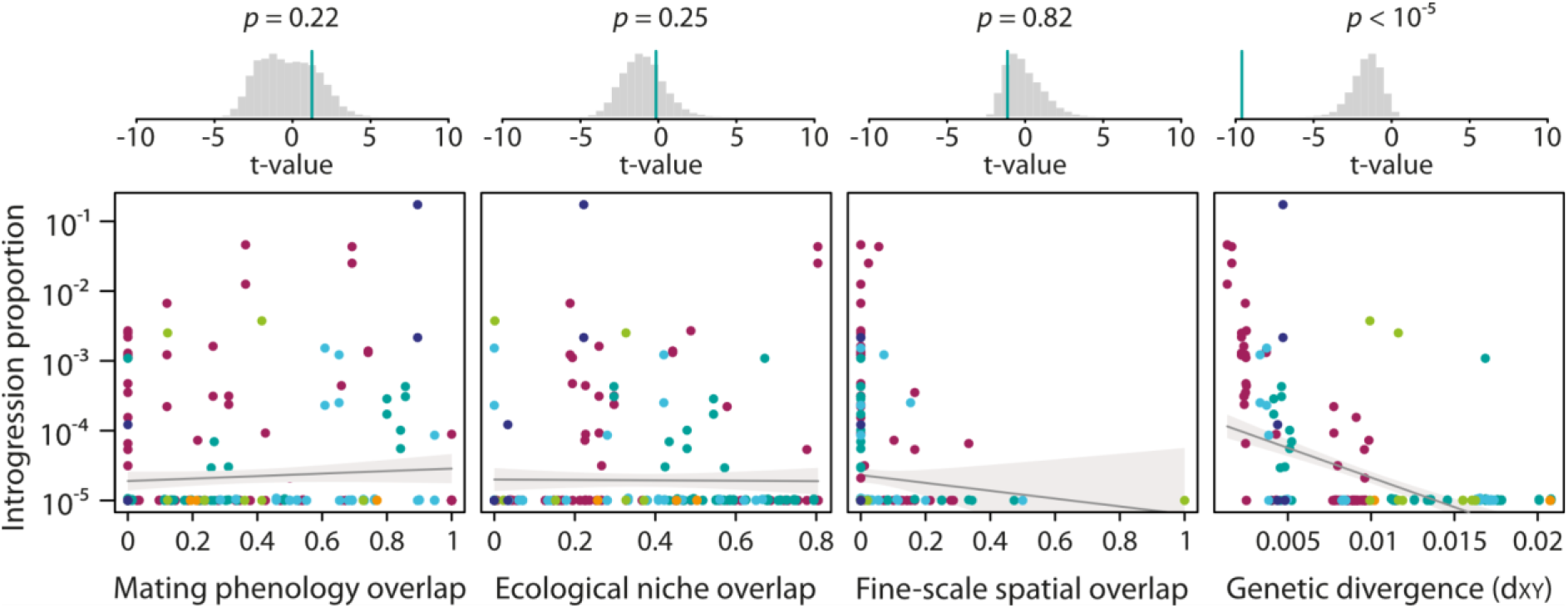
Relationship between introgression proportion and different pre-zygotic (mating phenology, ecological niche and fine-scale spatial overlap) and post-zygotic (divergence) reproductive barriers. Colors according to the genus, as in **Figure 1**. The predicted values and confidence intervals are represented by the grey line and area, respectively. Histograms at the top represent the distribution of simulated *t*-values under permutations. The green line is the observed *t*-value. The *p*-value is the proportion of simulated *t*-values falling beyond the observed one (one-tailed tests).

Why was there no association between prezygotic barriers and introgression? Prezygotic barriers were long believed to arise first and to play the major role in total reproductive isolation (*29–32*). This makes our result unexpected. One possibility is that we did not capture the most relevant barriers, such as chemical communication cues. However, our data instead suggest that the position of species along the speciation continuum may be more important. Many of the species pairs showing introgression were relatively old and often not sister taxa. This raises the possibility that prezygotic barriers may have eroded over time. Such erosion could occur through selection if these barriers are costly (as in *Phlox* flowers; or through drift if they become redundant or rarely expressed in allopatry. Resumed secondary contact after a long period in allopatry, which could be fueled for instance by anthropogenic disturbance (*33*), could then re-establish opportunities for hybridization between diverged species. Consistent with this interpretation, the hybridizing species we identified fall exactly within the “grey zone” of divergence where gene flow is still expected (*34, 35*). Our results mirror a large comparative study in plants, which similarly revealed a strong effect of genetic divergence but a low impact of distribution overlap on introgression (*11*).

Is introgression generally overlooked, or is it especially high in ants? Several ant-specific features, such as the absence of fertility cost for hybrid workers and haploid males (*36*, but see *37*), cheating genotypes that preferentially develop into reproductives (*38*) and the presence of lineage with hybrid-only workers (social hybridogenesis; *39*), could in theory make them specifically prone to hybridization (*15, 40*). However, these mechanisms would generate F_1_ hybrids instead of introgression. By contrast, almost all of the hybrids we identified had admixture proportions much lower than 50%, which indicates backcrosses and introgression. In addition, the introgressed proportion (species-wide proportion of the genome derived from other species) was low in the majority of species, ranging from 0.001 % in *Formica lemani* to 17.3% in *Tetramorium immigrans* (Figure 1D). Our results thus suggest that the high prevalence of introgression we observe does not reflect ant-specific biology, but rather the increased resolution of our approach. Instead of the common idea that a few species hybridize extensively (e.g., *15, 17*), our data is painting a different picture: many species hybridize at low levels, and this pervasive but weak introgression has largely gone unnoticed.

In summary, our assemblage-wide genomic survey reveals that introgression is widespread, but typically subtle. Rather than a few species hybridizing extensively, we find that many species exchange genetic material at low levels, a pattern that has likely gone undetected due to limited sampling and resolution. Despite its low magnitude, introgression makes a significant contribution to standing genetic variation and is primarily constrained by genetic divergence, consistent with postzygotic barriers playing the main role in limiting gene flow at this stage of divergence.

These results suggest that introgression may be a common feature of natural communities, with implications for how species boundaries are defined and maintained. More broadly, they highlight the importance of community-scale genomic approaches for quantifying evolutionary processes in the wild. Extending such approaches across diverse taxa will be essential to determining how pervasive introgression is across the Tree of Life and to understand its role in shaping adaptation and diversification.

## Materials and Methods

### Study area and sampling

We collected ants in the canton de Vaud, a 3212 km^2^ area in Switzerland which features a wide range of habitats and altitudes. The ants originate from two sampling efforts, conducted during the summer 2019. First, a structured sampling at 44 sites scattered across the study area and covering all habitat types (see (*41*) for details). Each site consisted of one square kilometer in which ants were searched in 25 circular plots of 4 meters of diameter, and along two kilometers of transects. The second source of samples was a citizen-science sampling campaign during which members of the general public collected samples anywhere within the study area (*42*). In both cases, about 10 workers were collected per colony and stored in 70% EtOH until DNA extraction. All samples were identified morphologically to the species level by expert taxonomists following (*43*).

We focused on all seven genera in which more than one species had at least 10 samples (*Camponotus, Formica Lasius, Myrmica, Tapinoma, Temnothorax*, and *Tetramorium*). We genotyped all individuals from all species for which we obtained fewer than 200 samples, and down-sampled the other species to 200 - 250 samples, aiming to maximize geographic coverage. The total number of samples selected was 5144: 174 *Camponotus*, 2094 *Formica*, 1171 *Lasius*, 758 *Myrmica*, 233 *Tapinoma*, 258 *Temnothorax*, 456 *Tetramorium*.

### Wet lab and sequencing

We randomly selected one worker per colony for genetic analyses. DNA was extracted from three legs. Individuals were grouped by species on the extraction plates, such that potential contaminations between adjacent wells would not artificially inflate gene flow between species. DNA extraction and COI barcoding of *Camponotus, Lasius, Myrmica, Temnothorax*, and *Tapinoma* was performed by the Canadian Center for DNA Barcoding. A fragment of the COI mitochondrial gene was amplified using primers LepF1 and LepR1 (*44*). The specific protocols used for extraction, PCR amplification and sequencing are available at https://ccdb.ca/resources (last checked 24.11.2025). DNA extraction and COI barcoding of *Formica* and *Tetramorium* was performed by AllGenetics & Biology SL, Spain. The same primers were used for sequencing.

We generated RAD sequencing data for all individuals using the protocol described in Brelsford *et al*. (*45*). Briefly, we digested genomic DNA with restriction enzymes EcoRI and MseI, ligated custom barcoded adapters, amplified the resulting fragments in 16 PCR cycles and selected fragments ranging between 280 and 430 bp using 2% agarose cassettes on a BluePippin (Sage Science). The resulting libraries were sequenced on the Illumina HiSeq 2500 platform (single end, 150 bases), on a total of 55 lanes (*Camponotus*: 2; *Formica*: 22; *Lasius*: 12; *Myrmica*: 8; *Tapinoma*: 3; *Temnothorax*: 3; *Tetramorium*: 5).

### SNP calling

We built SNPs using *stacks* v2.3e (*46*). We first demultiplexed the reads using the *process_radtags* command with the options *-c -q -r -t 143*. We then used the competitive mapping approach of our decontamination pipeline (see next section), and used the retained reads for SNP calling. We then mapped the reads onto one published genome assembly per genus: *Camponotus fallax* (accession number GCF_003227725.1); *Formica selysi* (GCA_009859135.1); *Lasius niger* (GCA_041902855.1); *Myrmica rubra* (GCA_048181765.1); *Tapinoma erraticum* (*47*); *Temnothorax unifasciatus* (GCA_048541725.1) and *Tetramorium immigrans* (GCA_011636585.1). We mapped reads using the *mem* algorithm of *bwa* version 0.7.17 (*48*) and used *samtools* version 1.4 (*49*) to convert mapping files into bam format and to generate summary statistics. We then built loci using *gstacks*, enabling the options *--phasing-donprunhets* and *--ignore-pe-reads*. Finally, we ran *populations* with the option *--write-single-snp* to retain only one SNP per RAD locus. We also ran *populations* with options *--vcf-all* and *-r 0.05* to create a genomic VCF (gVCF) file for nucleotide diversity estimations.

We filtered the data in *vcftools* 0.1.14 (*50*). We retained genotype calls with a minimum depth of 8, and loci for a minor allele count of two and presence in at least 75% of individuals. We discarded all individuals which had over 50% of missing data. We generated datasets for subsets of individuals for subsequent analyses by filtering with the same parameters. All downstream analyses were carried out in R 4.1.1 (*51*).

### Avoiding potential contamination

Cross-contamination between samples belonging to different species can mimic gene flow and introgression. To detect and remove them, we used our previously developed pipeline (*52*), which involves two steps. First, to remove potential contaminants from different ant taxa, we mapped the raw RADseq reads to a concatenated file containing the assemblies used for each of the seven genera (see species and accession numbers above). We only retained reads that mapped to the assembly of the expected genus for SNP calling (see above).

The second step is based on allelic depth ratio (a.k.a. allelic imbalance), which we define here for each heterozygous site as the proportion of reads supporting the major allele. Allelic depth ratio (hereafter ADR) thus ranges between 0.5 (even) towards 1 (highly skewed). We estimated the expected distribution of ADR ratio across all observed heterozygous sites in each vcf. Any individual whose median ADR was above the 95% percentile of this expected distribution was considered contaminated and discarded. In addition, any site at which ADR was higher than the 75% percentile of the expected distribution was corrected by removing the allele with the lowest depth. All downstream analyses were then run on this filtered dataset.

### Corroborating morphological species identification with genetic data

Previous work in ants showed that, while relatively accurate, morphological identification can result in misidentification of a small proportion of individuals (*53*). In order to corroborate morphological identifications, we built a phylogeny on the COI sequence and conducted non-metric Multidimensional Scaling (NMDS) analyses and unsupervised *admixture* analyses (*54*) on the RAD data. The COI data was aligned with *mafft* 7.525 (*55*) and the phylogeny was then built with *iqtree2* 2.2.2.7 (*56*). The NMDS analysis was run with the *isoMDS* function of the R package *MASS* (*57*) on the euclidean genetic distances between individuals. All analyses were run for each genus separately, as both the rate of intergeneric hybridization and the proportion of RADtags showing enough homology across all genera to be informative are expected to be very low. Because admixture and related approaches perform poorly with strong sample size biases, we excluded species represented by fewer than 10 individuals from subsequent analyses.

### Estimating introgression rate

Measuring the rate of introgression as the number of hybrids found between pairs of species would be subject to bias, because it requires setting a cutoff of ancestry contribution from a second species above which an individual is considered hybrid. However, low ancestry proportions of a second species can also represent errors, for example due to missing data.

We thus measured hybridization rate between all pairs of species from the same genus as the genome proportion from all individuals belonging to one species that is derived from the other species. To this end, we first identified the five most pure-species individuals from each species with the lowest missing data. We used them as references in a supervised admixture analysis, setting k as the number of species analyzed within the genus. We ran admixture in 20 replicates and selected the best run (i.e. with lowest cross validation error). We assigned each individual to the species from which they had the largest ancestry proportion. We then computed hybridization rate asymmetrically, from species A into species B, as the sum of A-derived ancestry proportions of all individuals of species B, and vice versa from species B into species A, as the sum of B-derived ancestry proportions of all individuals assigned to species A.

We then estimated the proportion of species showing some signs of introgression as the number of species with nonzero introgression values, divided by the total number of species found in the area, including those not genotyped. Our estimation of the proportion of hybridizing species is thus a conservative, minimum estimate. It is however realistic, as species that are unique representatives of their genus most likely lack hybridization opportunities.

### Contributions of introgression to genetic diversity

We estimated the contribution of introgressed individuals to nucleotide diversity (π) within each of the species that engaged in introgression. For each species, we first generated a list of pure individuals (i.e. individuals showing no signs of introgression), and a list of all individuals, including introgressed ones. We then used the gVCF file including all sequenced positions generated by stacks. For both lists of individuals, we generated a subset of the gVCF containing individuals present in the list with *vcftools* (*50*). We estimated nucleotide diversity (π) on that gVCF using *pixy* (*58*), with a window size of 100 kb. We then estimated average genome-wide π by summing all nucleotide differences for all windows and dividing by all nucleotide comparisons across all windows. In *Formica*, we excluded the entire chromosome 3, as it is a trans-species supergene under balancing selection (*59*), which would bias the estimate by inflating π in heterozygotes.

We then estimated what increase in population size would be responsible for the same amount of increase in π as our observed value for introgression. We used the regression by (*22*), who found that a 10-fold increase in census population size resulted in a 13% increase in π, a pattern consistent across taxa and population sizes. Because both Nc and π are on a log10 scale in the study by Buffalo (*22*), we estimated the census size (N_c_) increase that would be equivalent to our estimated π increase as:

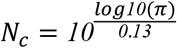

where π is the average ratio of π with / without hybrids.

### Factors associated with introgression rate

We asked whether classical reproductive barriers were associated with introgression rate. We first measured three of the main prezygotic barriers known in ants and in general (reviewed in *60*). The first is the overlap in mating phenology, which reflects hybridization opportunities between species. The second is the overlap in ecological niche, defined from 59 environmental variables. The third is fine-scale local co-occurrence, which captures both large-scale distribution and fine-scale microhabitat use. We describe how these variables were estimated below. The prediction is that all three variables should be positively associated with introgression rate.

Finally, we assessed the effect of genetic divergence (d_XY_), as it is supposed to be directly linked to genetic incompatibilities (*61*), which is the main postzygotic barrier. It is thus predicted to be negatively associated with introgression rate.

We estimated the relationship between the log_10_-transformed introgression rate and each of these potential barriers independently. Because of the structure of our data (species nested within genus, and each species acting both as “donor” and “receiver” of introgression), we tested the significance of the relationships using permutations in a Mantel test-like manner. We permuted the introgression rate (our response variable) in our data matrices symmetrically, within each genus: when rows *i* and *j* were permuted, so were columns *i* and *j*. We extracted the observed *t*-value of the linear model, and compared it to the distribution of the *t*-values of the linear models of datasets permuted as described, across 10’000 replicates. The *p*-value of the model was estimated as the proportion of the *t*-values in the randomized datasets that were higher (for phenology overlap, ecological niche overlap and fine-scale spatial overlap) or lower (d_xy_) than the observed *t*-value. All analyses were run in R 4.1.1 (*51*).

### Estimating mating phenology overlap

Phenology of each species was estimated based on the earliest and latest swarming dates reported by (*43*). Those are largely based on observations made in Germany. Swarming dates might vary with latitude and altitude, but our study area is located close to Germany and experiences a similar climate. In addition, while phenology of the whole community might be impacted by local climatic variation, the relative phenology of different species is unlikely to change drastically.

We measured seasonal phenology overlap as the Jaccard index of the ordinal dates (i.e. ranging from 1 to 365 where 1 = January 1^st^ and 365 = December 31^st^), computed as

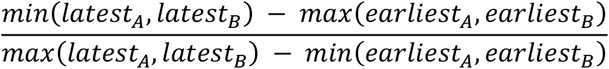

where *latest*_*A*_ and *latest*_*B*_ are the latest, and *earliest*_*A*_ and *earliest*_*B*_ are the earliest swarming days of the year for species A and B, respectively. For the few species with two swarming periods, both were taken into account.

We followed the same approach with daily phenology, computing the Jaccard index of the number of minutes from midnight. We obtained daily phenology from the same source (*43*). In the few cases where reported swarming time was relative to sunrise or sunset, we converted it into minutes from midnight using the time of sunrise and sunset in Lausanne, which is central in our study location, on the mean swarming date of the species.

Finally, we also computed the combined phenology overlap by multiplying seasonal and daily phenology. Because the results for these three measures were qualitatively similar and seasonal phenology was available for more species, we present results from daily and combined phenology overlap in Figure S9.

### Estimating ecological niche overlap

We estimated ecological niche overlap between all species using the framework developed by (*62*). This approach quantifies niche overlap by projecting species occurrences into a common multivariate environmental space (PCA-env), estimating smoothed occurrence densities, and comparing them while accounting for the availability of environmental conditions in the study area. Ecological niches were estimated using only occurrences of genetically pure individuals (assignment probability to their main cluster > 99.9%) to avoid biases due to admixture. We characterized the environmental space using 59 ecological variables relevant to ant ecology, including climate, land use, urbanization, vegetation productivity (NDVI) and canopy height.

Climatic variables at 25 m x 25 m resolution, including precipitation, temperature, growing season and bioclimatic variables, were extracted from the CHClim25 dataset (*63*). The aspect and slope were calculated from a Digital Elevation Model using the *terrain* function from the R package *raster* (*64*). Land-use variables quantified the proportion of each pixel covered by major habitat categories (crops, forests, edges, and urban habitats). To account for ant mobility (*43*), the proportion of each land-use category was also calculated in a 25 m and 200 m neighborhood around each pixel using the function *focal* from *raster*. Anthropogenic pressure was characterized by the length of roads and the perimeter of buildings in each pixel using information derived from OpenStreetMaps. Soil conditions, continentality and light were captured using plant Ecological Indicator Values in each pixel (*65*). Vegetation productivity was quantified using average canopy height and the mean Normalized Difference Vegetation Index (NDVI) values by pixel, with NDVI averaged over the period 2010-2020 from 206 LANDSAT 5 and LANDSAT 8 images processed via Google Earth Engine. In total, we assembled an initial set of 59 predictors with a resolution of 25 m x 25 m over the study area. Variable pre-processing was performed in R 4.1.1 and QGIS 3.16.13. We then built an environmental Principal Component Analysis (PCA) on all the pixels of the study area using the full set of 59 predictor variables.

Because of potential biases in the citizen sampling campaign (such as preferential sampling in some habitat types or detectability differences between species), we corrected citizen-science occurrences using an environmental down-sampling procedure. We first defined eight environmental types based on the first two principal components of the environmental PCA (Figure S10). For each species, we then tested whether the proportion of observations across these environmental types differed significantly between citizen and structured sampling using a χ^2^ test. For species where there was no significant difference, we used all observations from both citizen and structured sampling together as occurrences. For the 37.5% of species where the χ^2^ test was significant (with *p* < 0.05), we down-sampled citizen-science occurrences so that their proportions across the eight environmental types matched that of the structured sampling.

We then quantified the ecological niche overlap of each species pair using the approach of (*62*) implemented in the R package *ecospat* v3.4 (*66*). To reduce computation time, we estimated the overlap on a random subsample of 10’000 environmental points across the area. The overlap between the niches of each species was measured using Schoener’s D statistic, where 0 indicates no overlap and 1 complete overlap.

### Estimating fine-scale spatial overlap

In order to test whether co-occurring species were more likely to introgress, we estimated fine-scale spatial overlap between species. For this, we leveraged the data from the structured sampling, which recorded which species were found in the same 4-meter circles. We estimated co-occurrence directionally: co-occurrence of A with B was measured as the number of plots where species A and B co-occurred divided by the number of plots where A occurred.

### Estimating genetic divergence between species

We tested whether genetic distance between species had an influence on admixture proportion. We used nucleotide divergence (d_XY_) as a proxy for genetic divergence. We estimated d_XY_ from the same gVCF containing only non-hybrid individuals as for π using pixy (*58*), with a window size of 100 kb. We then estimated average genome-wide d_XY_ by summing all nucleotide differences between species and dividing by all nucleotide comparisons. In *Formica*, as above, we excluded the entire chromosome 3, as it is a trans-species supergene under balancing selection (*59*).

In order to be able to compare our divergence values with previous studies (*34, 35*), we also estimated net nucleotide divergence (d_A_) for each window following eq. 25 in (*67*) as

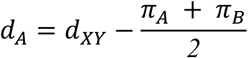

where d_XY_ is averaged over the whole genome and π_A_ and π_B_ are average genome-wide π values for species A and B, respectively.

## Acknowledgements

We are thankful to the 608 people who collected ants as part of the “Opération fourmis” citizen-science inventory. We thank Tim Szewczyk for the structured sampling; Anne Freitag, Aline Dépraz and Amaury Avril for coordinating the citizen-science sampling campaign; Anne Freitag, Sebastian Salata, Christophe Galkowski and Claude Lebas for ant identification; Marc Bastardot, Marjorie Labédan, Christine La Mendola, Heather Ryder and Simon Vogel for help with processing the samples; Claude Lebas and Hugo Darras for providing reference samples of *Tapinoma*.

This study was supported by funding from the University of Lausanne and the Herbette foundation. GL is supported by a Postdoc.Mobility fellowship from the Swiss National Science Foundation (grant number P500PB_222111).

## Author contributions

Designed the study: GL, TS. Obtained the samples: TS. Wetlab: GL, LR. Analyses: GL, OB, MT, KJ, LR, with input from TS, JG. Writing: GL, with input from TS and all authors.

## Data availability

All demultiplexed RAD reads are available on NCBI: PRJNA1458683 (*Camponotus*), PRJNA1466771 (*Formica*), PRJNA1176511 (*Lasius*), PRJNA1188312 (*Myrmica*), PRJNA1476096 (*Tapinoma*), PRJNA1476131 (*Temnothorax*), PRJNA1476143 (*Tetramroium*). All code is available on GitHub: https://github.com/glavanc1/gene_flow_ants.

