## Supplementary material for "Widespread but cryptic introgression shapes genetic diversity in natural populations"

For the manuscript entitled "**Widespread but cryptic introgression shapes genetic diversity in natural populations**"

### Corroborating morphological species identification

We used an integrative approach including combined information from morphological identification, COI barcoding and RAD sequencing  to assign each individual to a species for each genus, as described in (*53*). Briefly, taxonomists identified as many individuals as possible to the species level following the taxonomy of (*43*). Two species groups could not be identified morphologically to the species level: *Tapinoma nigerrimum* grp. and *Formica lugubris/paralugubris*. We used the genetic information instead.

The genetic species identification was largely congruent with morphological identificationAfter excluding all individuals with more than 75% missing data and all species with fewer than 10 individuals with sufficient data, we retained three *Camponotus* species (134 individuals): *fallax, herculeanus, ligniperda*; 12 *Formica* species (1640 individuals): *cunicularia, exsecta, fusca, lemani, lugubris, paralugubris, polyctena, pratensis, pressilabris, rufa, rufibarbis, sanguinea*; eight *Lasius* species (952): *L. brunneus*, *L. emarginatus*, *L. flavus, L. fuliginosus*, *L. mixtus*, *L. niger*, *L. paralienus* and *L. platythorax*; seven *Myrmica* species (652 individuals): *lobulicornis, rubra, ruginodis, sabuleti, scabrinodis, schencki, specioides*; 3 *Tapinoma* species (198 individuals): *erraticum, magnum, subboreale*; four *Temnothorax* species (201 individuals): *affinis, nigriceps, nylanderi, unifasciatus*; three *Tetramorium* species (349 individuals): *caespitum, immigrans, impurum*. In total, we retained 4126 individuals in 40 species from seven genera.

*Camponotus*

Morphological identification revealed the presence of five species (*C. aethiops, C. fallax, C. herculeanus, C. ligniperda, C. piceus*). The only sample of *C. aethiops* did not pass genetic data filtering. The genetic analyses confirmed the presence of four clusters, each containing individuals identified morphologically as one species, with the exception of two individuals clustering with *C. herculeanus* that were identified as *C. ligniperda* (Figure S1).

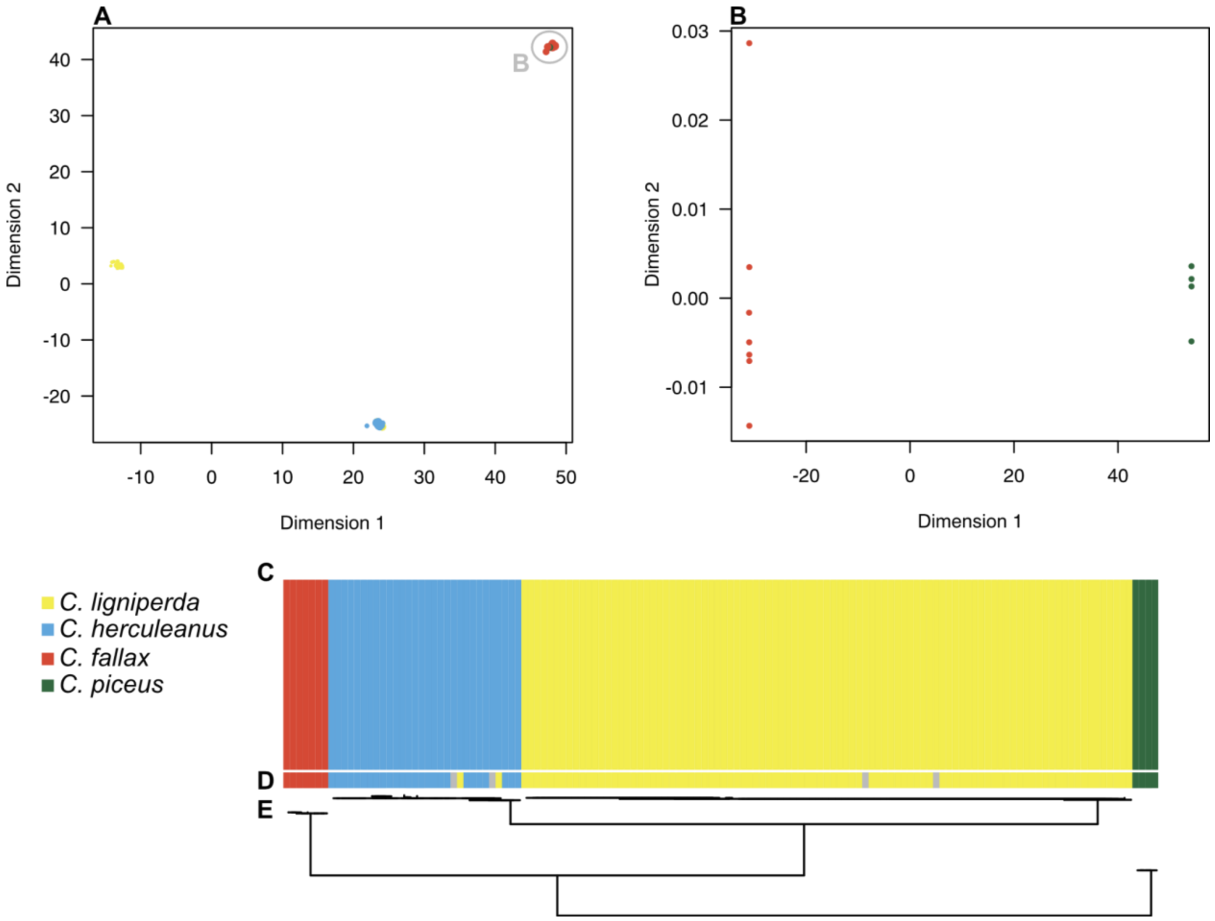


**Figure S1**: Genetic identification of *Camponotus* samples. **A**: NMDS plot of all samples. The light gray circle indicates the cluster containing a mix of samples identified morphologically as *C. fallax* and *C. piceus*. **B**: NMDS plot of the subset of samples identified morphologically as *C. fallax* and *C. piceus*. Each point is one individual, coloured according to its morphological species identification. Points in grey designate individuals for which morphological identification was unsure. **C**: Results of admixture with k = 4. **D**: morphological identification. **E**: phylogeny of the COI gene.

*Formica*

The NMDS separated all described subgenera (**Figure S2**) but did not provide enough resolution to separate all species, which prompted us to split the data into subsets. Analyses of subsets then enabled identification of all individuals. We  ran admixture analyses separately for the *Formica s. str* group and other species of the genus (Figure **S2H** and **I**). The COI phylogeny revealed a lack of polymorphism in the analyzed gene portion for some closely related species, grouping *F. rufa* and *F. polyctena*, as well as *F. fusca* and *F. lemani*, both of which were otherwise morphologically and genetically distinct. In addition, *F. lugubris*, *F. paralugubris* and *F. pratensis* were paraphyletic on the COI phylogeny. There were six main haplotypes, each corresponding to a single cluster (with the exception of some hybrids). One haplotype corresponded to most individuals identified morphologically as *F. pratensis* and was nested within the others. The other five contained most individuals identified as *F. lugubris* / *paralugubris* and were split in two groups by admixture and the NMDS, each haplotype containing only individuals belonging to one of these clusters, which is indicative of incomplete lineage sorting. The cluster of the *F. lugubris* / *paralugubris* group encompassing the reference individuals of *F. aquilonia* (which is represented in dark magenta in

**Figure S2H**) was inferred to be *F. paralugubris* because the latter is hypothesized to be a hybrid species between the former and *F. lugubris* (*68*).

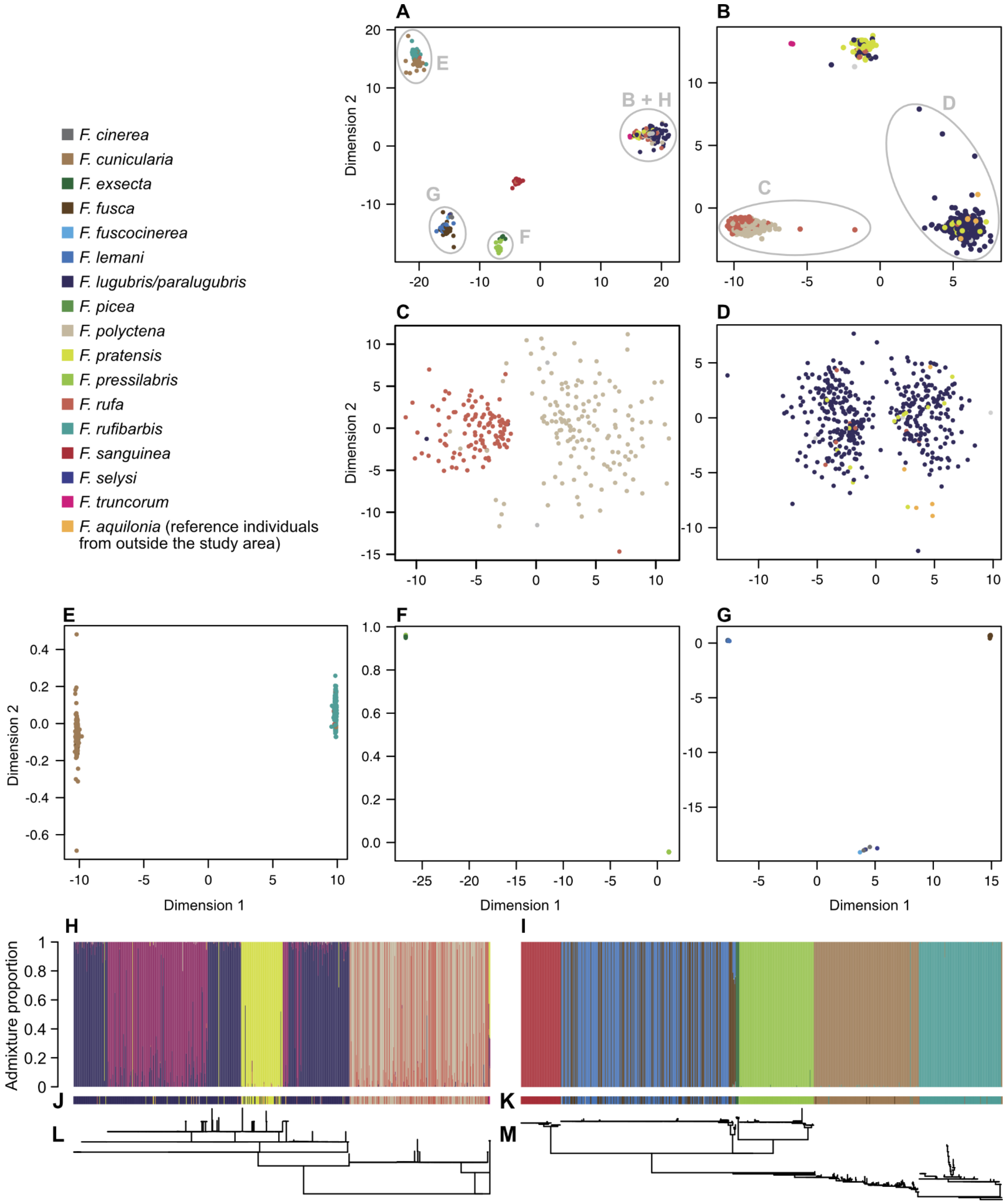


**Figure S2**: Genetic identification of 1659 *Formica* samples. **A**: NMDS plot of all samples. Light gray circles indicate subsets of individuals for which additional analyses were run, and corresponding letters indicate panels where those analyses are presented. **B**: NMDS plot of samples of the subgenus *Formica s. str.* **C**: NMDS plot of the *F. rufa* / *F. polyctena* group. **D**: NMDS plot of the *F. lugubris* / *F. paralugubris* group. **E**: NMDS plot of the *F. cunicularia* / *F. rufibarbis* group. **F**: NMDS plot of the *F. exsecta* / *F. pressilabris* group. **G**: NMDS plot of the group encompassing *F. cinerea*, *F. fuscocinerea*, *F. lemani* and *F. selysi*. **H** and **I**: Results of admixture for the *Formica s. str.* group with k = 5, and for the other species with k = 7, respectively. **J** and **K**: Morphological identification of individuals presented in **H** and **I**, respectively. **L** and **M**: phylogeny of the COI gene for all individuals presented in **H** and **I**, respectively. Note that both phylogenies have different scales. Forty-four individuals of *F. rufibarbis* showed an apparently very divergent COI haplotype, indicative of a nuclear mitochondrial transfer (NUMT), and were excluded from panels **I**, **K** and **M**. The NMDS plots showed no indication of a different nuclear lineage. Note that the assignment of ambiguous individuals (e.g. those with a value close to 0 on the x axis in **C** and **D**) to one or the other cluster has no effect on downstream inference of introgression rate. This is because introgression rates were estimated from a second admixture analysis excluding rare species, which this time was "supervised" (i.e. taking the purest individuals as references to estimate allele frequencies to define clusters – see methods). Ambiguous individuals were thus not used to define clusters, regardless which species they belong to.

*Lasius*

Genetic species identification allowed un-ambiguous assignment of individuals for nine out of 12 species: *L. brunneus*, *L. flavus, L. fuliginosus*, *L. mixtus*, *L. myops*, *L. emarginatus*, *L. niger*, *L. platythorax* and *L. paralienus*. Genetically similar species (with partial overlap in the latter on the NMDS including all samples) clearly separated in analyses based on species subsets (Figure S3). One species, *L. umbratus*, was represented by only one morphologically-identified individual, but this individual clustered genetically with *L. mixtus*, so we conclude that it was a morphological mis-identification. Because of their low sample size, the two remaining species (*L. alienus*, *L. neglectus*) appeared as admixed between the major species on the admixture analysis. However, they were separated on the NMDS and had distinct COI haplotypes, and we conclude that they were correctly identified. We discuss the taxonomic implications of our results in the *Lasius* genus in more detail in (*52*).

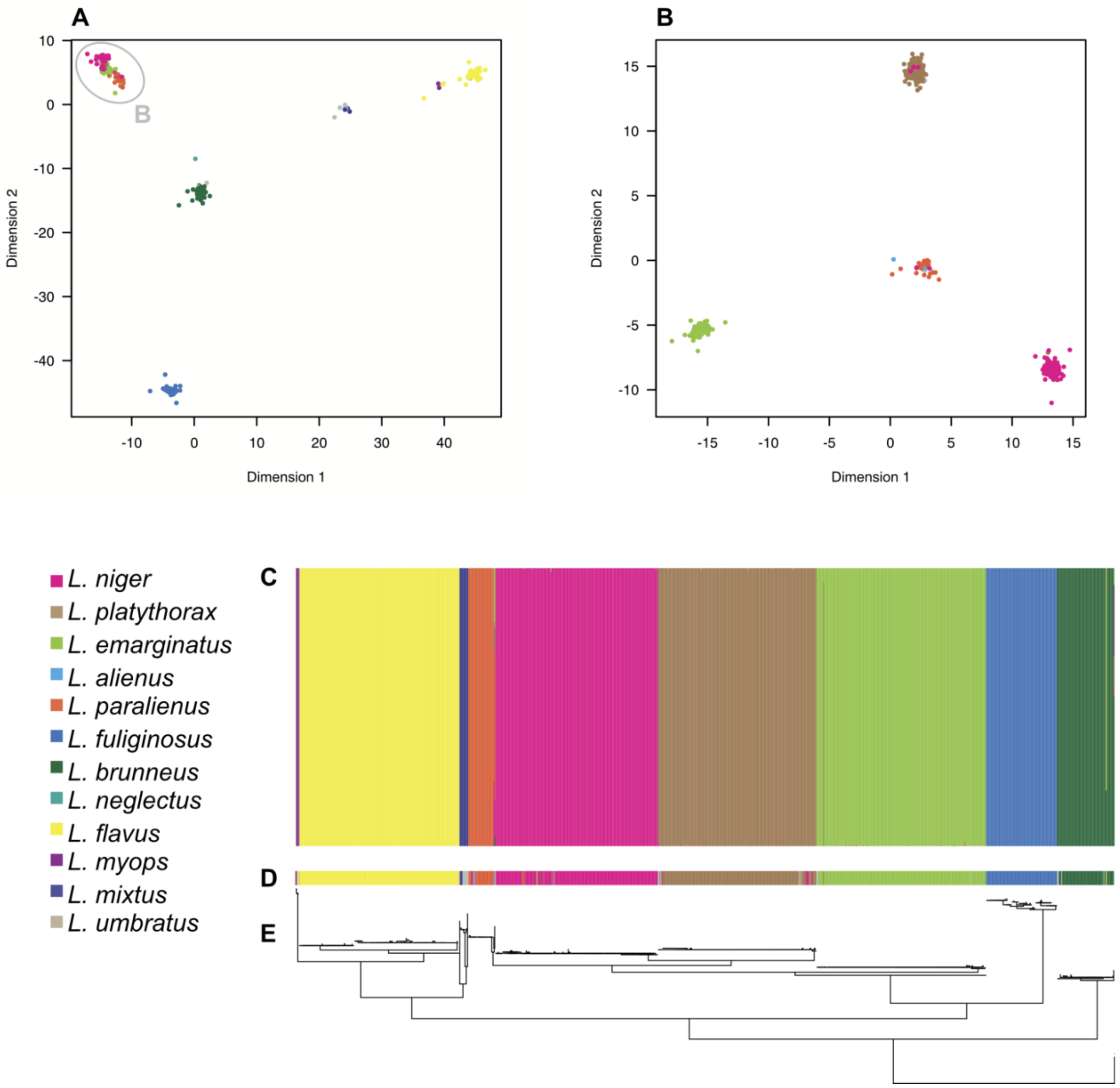


**Figure S3**: Genetic identification of *Lasius* samples. **A**: NMDS plot of all samples. The light gray circle indicates the cluster of samples of the *L. niger* group. **B**: NMDS plot of the subset of samples of the *L. niger* group. Each point is one individual, coloured according to its morphological species identification. Points in grey designate individuals for which morphological identification was unsure. **C**: Results of admixture with k = 9. **D**: morphological identification. **E**: phylogeny of the COI gene.

*Myrmica*

The analysis of this dataset was the subject of a previous study (*53*). Briefly, five species were clearly identified by all analyses: *M. rubra*, *M. ruginodis*, *M. sabuleti*, *M. specioides* and *M. sulcinodis*. A sixth species, *M. scabrinodis*, showed two paraphyletic mitochondrial haplotypes. However, we have shown previously that they do not reflect a cryptic species and are likely the result of incomplete lineage sorting (*53*). Individuals identified as *M. schencki*, *M. lobulicornis* and *M. lobicornis* clustered together in the admixture analysis but showed different mitochondrial haplotypes (although with very shallow divergence in the latter two). Four other species (*M. curvithorax*, *M. gallienii*, *M. rugulosa* and *M. vandeli*) were represented by very few samples and appeared admixed on the admixture analysis but had separate mtDNA haplotypes and appeared distinct from other lineage on the NMDS.

The only sample morphologically identified as the last species, *M. lonae*, appeared identical to *M. sabuleti*.

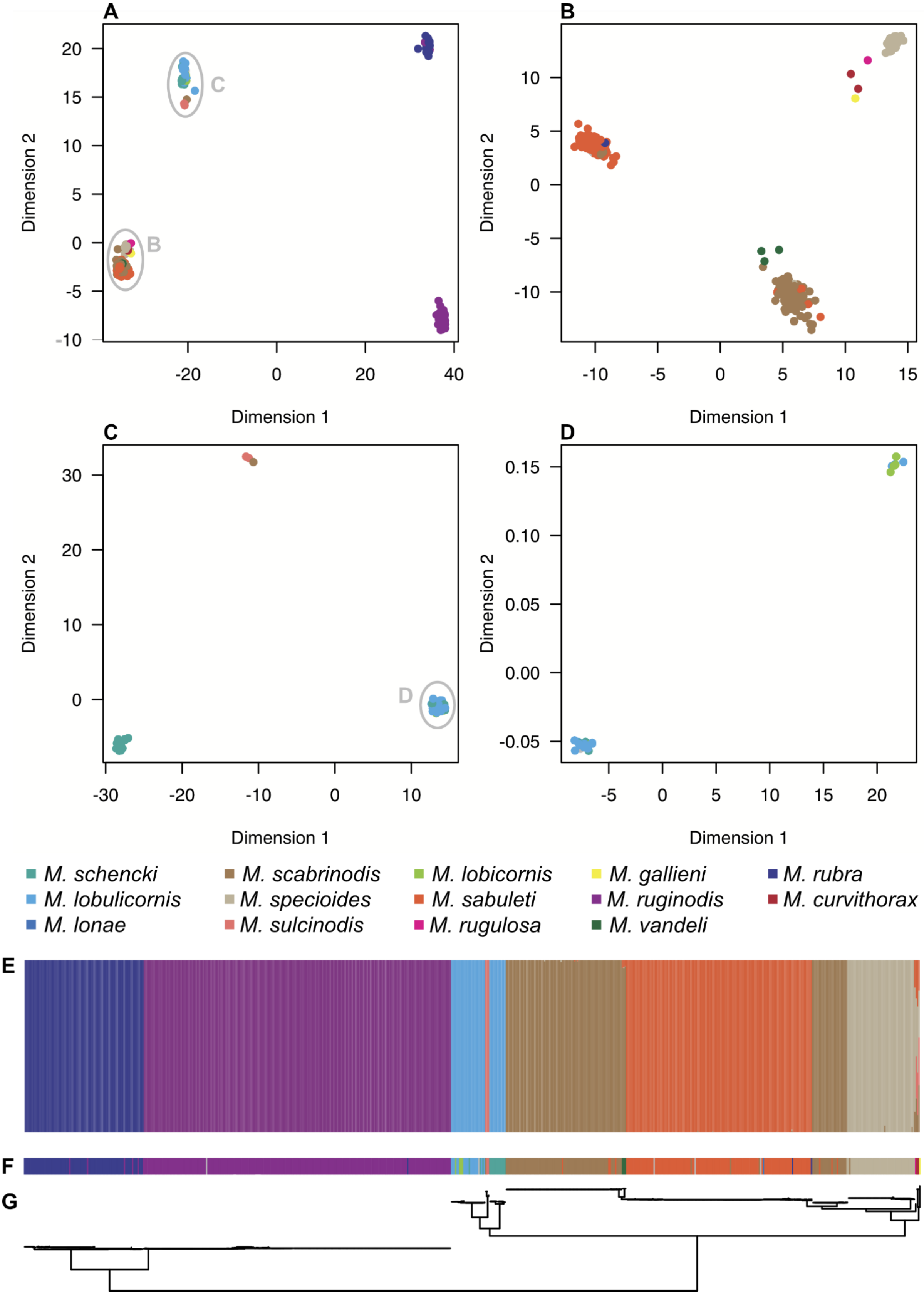

**Figure S4**: Genetic identification of *Myrmica* samples. **A**: NMDS plot of all samples. The light gray circles indicate clusters of samples reanalyzed separately. Each point is one individual, coloured according to its morphological species identification. Points in grey designate individuals for which morphological identification was unsure. **B**: NMDS plot of the subset of samples of the *M. sabuleti* group. **C**: NMDS plot of the subset of samples of the *M. schencki/lobulicornis/lobicornis* group **D**: NMDS plot of the subset of samples of the *M. lobulicornis/lobicornis* group. Note that artificial noise was added in **C** and **D** to improve display of otherwise overlapping points. **E**: Results of admixture with k = 7. **F**: morphological identification. **G**: phylogeny of the COI gene.

*Tapinoma*

Morphological identification revealed the presence of two species (*T. erraticum, T. subboreale*), plus one group of individuals belonging to the recently-described *T. nigerrimum* species group (*69*). One species from this group, *T. magnum*, was recorded in the study area for the first time two years before our inventory, and it has quickly become invasive. No other species from this group had ever been observed in the area.

Most individuals identified as *T. subboreale* formed a single cluster, along with some unidentified individuals and a few individuals identified as *T. erraticum*. The majority of individuals identified as *T. erraticum* formed a cluster, together with the majority of unidentified individuals, a few individuals identified as *T. subboreale* and one identified as *T. nigerrimum* grp. Finally, all but one individual identified as *T. nigerrimum* grp. clustered together, with one individual identified as *T. subboreale* and two unidentified. However, four individuals from this cluster presented a different mitochondrial haplotype (Figure S5).

In order to assess whether they could be two different species, and confirm the species identification, we added two samples from each of the four species of this group originating from within the native distribution of those species. The samples were provided and identified by Claude Lebas. We constructed RADseq libraries and called SNPs following the same protocol as described in the main text and at the same time as the samples from our study area. We then exported a phylip alignment using the *populations* module of *stacks* with the options *-r 0.5 -R 0.5 -p 2 --min-mac 2*, considering each individual as a separate population. We reconstructed the phylogeny with IQ-TREE version 2.2.2.7 (*56*).

We found that most individuals clustered with *T. magnum*, while the four individuals with a different mitochondrial haplotype clustered with *T. darioi*. This is the first record of *Tapinoma darioi* for Switzerland. Both *T. magnum* and *T. darioi* have the potential to become invasive, and monitoring the spread of these species will be crucial to estimate their ecological and economic impacts.

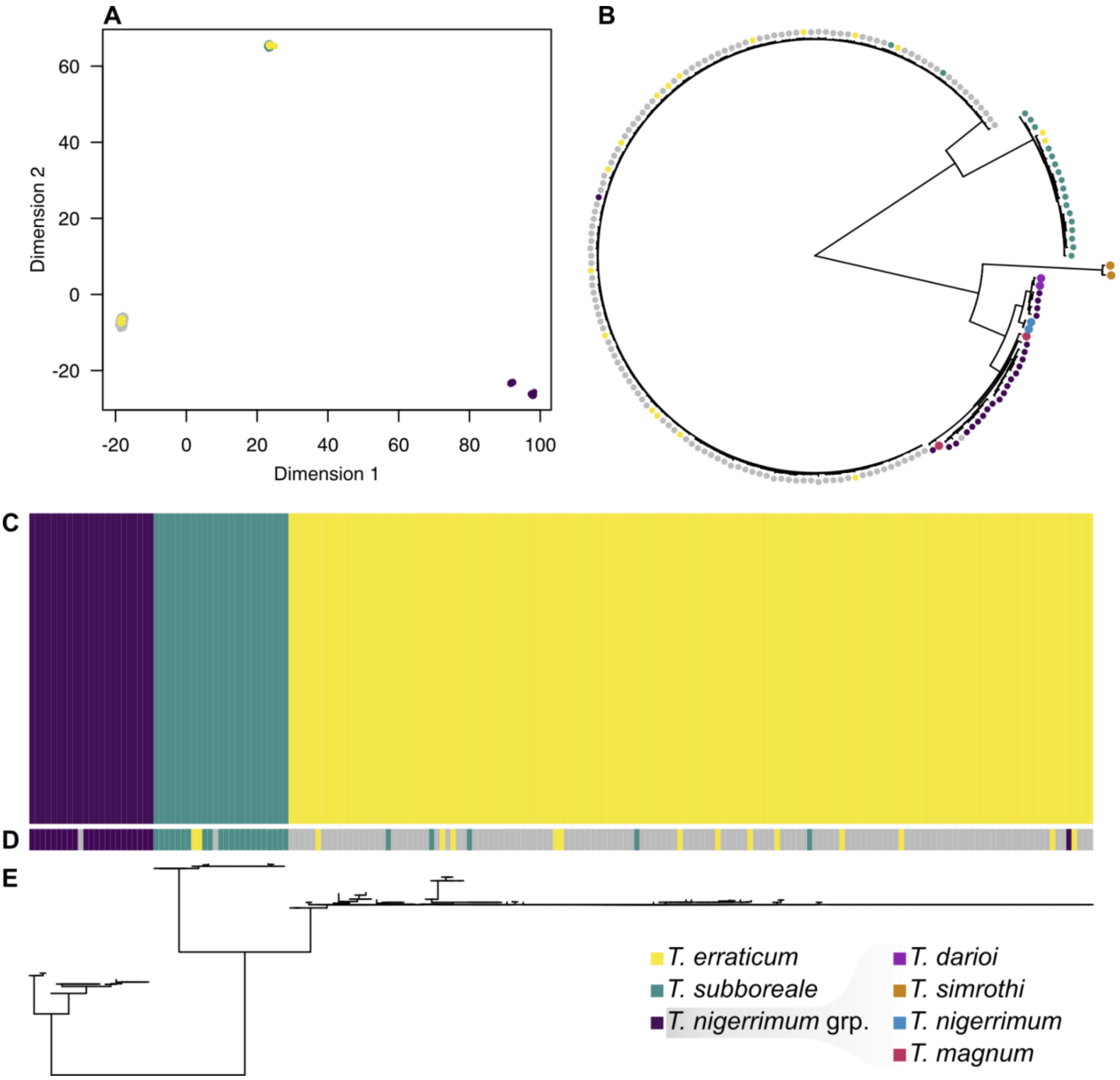


**Figure S5:** Genetic identification of *Tapinoma* samples. **A**: NMDS plot of all samples. Each point is one individual, colored according to its morphological species identification. Points in grey designate individuals for which morphological identification was unsure. **B**: phylogeny of all individuals including eight reference individuals (two per species of the *T. nigerrimum* group). Dot color represents morphological species identification, with larger dots for reference individuals. **C**: Results of admixture with k = 3. **D**: morphological identification. **E**: phylogeny of the COI gene.

*Temnothorax*

Two species (*T. affinis*, *T. interruptus*) showed a perfect match between morphological and genetic identification. Another species (*T. corticalis*) was represented by a single individual. The fact that it did not cluster with any other individual on the NMDS suggests that it was indeed *T. corticalis*, but amplification of its mtDNA failed, preventing us from genetically confirming the species. All individuals identified morphologically as *T. nylanderi* clustered together on the NMDS. Six individuals were identified morphologically as *T. parvulus*. Five of them clustered with *T. nylanderi*, and the last one clustered together with a second individual of unknown morphological species. Both had the same mitochondrial haplotype, which corresponded to published sequences of *T. parvulus*. Most individuals identified morphologically as *T. unifasciatus* clustered together on the NMDS and in admixture and had similar mtDNA, which did not correspond to any published sequence, but were closest to *T. cordieri*, a species known only from Corsica (*70*). However, one individual had the same mitochondrial haplotype as the only individual identified as *T. tuberum*, but the two differed in nuclear DNA. In addition, the only individual identified morphologically as *T. nigriceps* for which mtDNA amplification succeeded had a COI sequence matching that of published individuals of *T. tuberum*. We conclude that this individual is a true *T. tuberum*, and that the two other individuals are either a diverged lineage of *T. unifasciatus*, or another (maybe undescribed) species. Finally, COI amplification failed for all other individuals of *T. nigriceps*. This might suggest that there is a SNP in the sequence on which the primers anneal in this species. All individuals clustered together on the NMDS, with the exception of the ones discussed above.

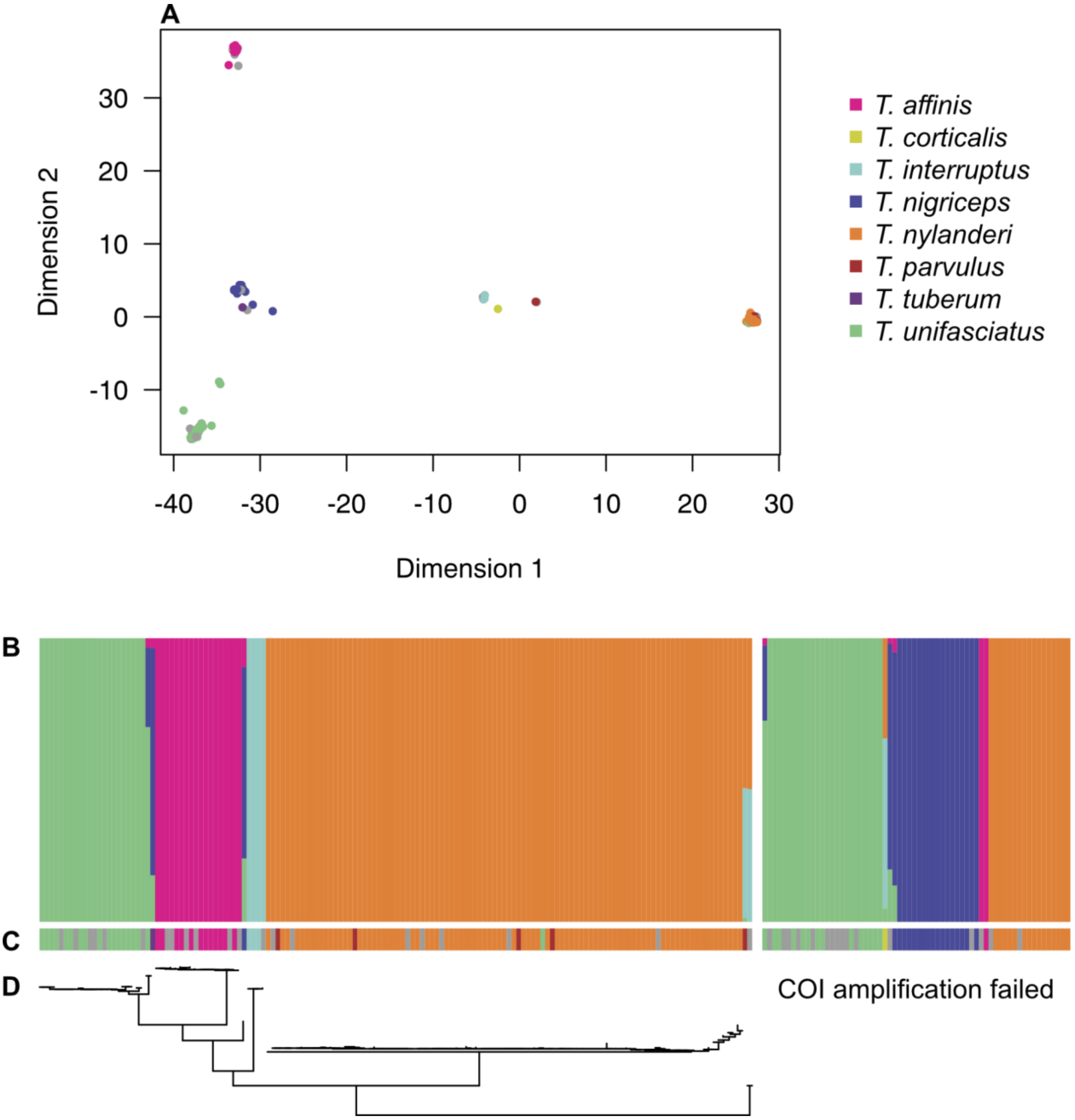


**Figure S6**: Genetic identification of *Temnothorax* samples. **A**: NMDS plot of all samples. Each point is one individual, coloured according to its morphological species identification. Points in grey designate individuals for which morphological identification was unsure. **B**: Results of admixture with k = 4. **C**: morphological identification. **D**: phylogeny of the COI gene.

### *Tetramorium*

Species of this genus are notoriously arduous to identify based on morphological criteria (Seifert 2018). Morphological specimen identification revealed the presence of five species (*T. alpestre, T. caespitum*, *T. immigrans*, *T. impurum*, plus one putative *T. semilaevae*). Our analyses revealed that most individuals belonging to *T. alpestre* and *T. impurum* in fact formed one single cluster (Figure S7). Their mitochondria were more similar to *T. impurum* than *T. alpestre*, so we conclude that they belong to *T. impurum*. The individual identified as T. semilaevae was in fact *T. caespitum*. Finally, the mitochondrial polymorphism in *T. caespitum* (already described in (*71*)) did not reflect substructure at the nuclear level (Figure S7C).

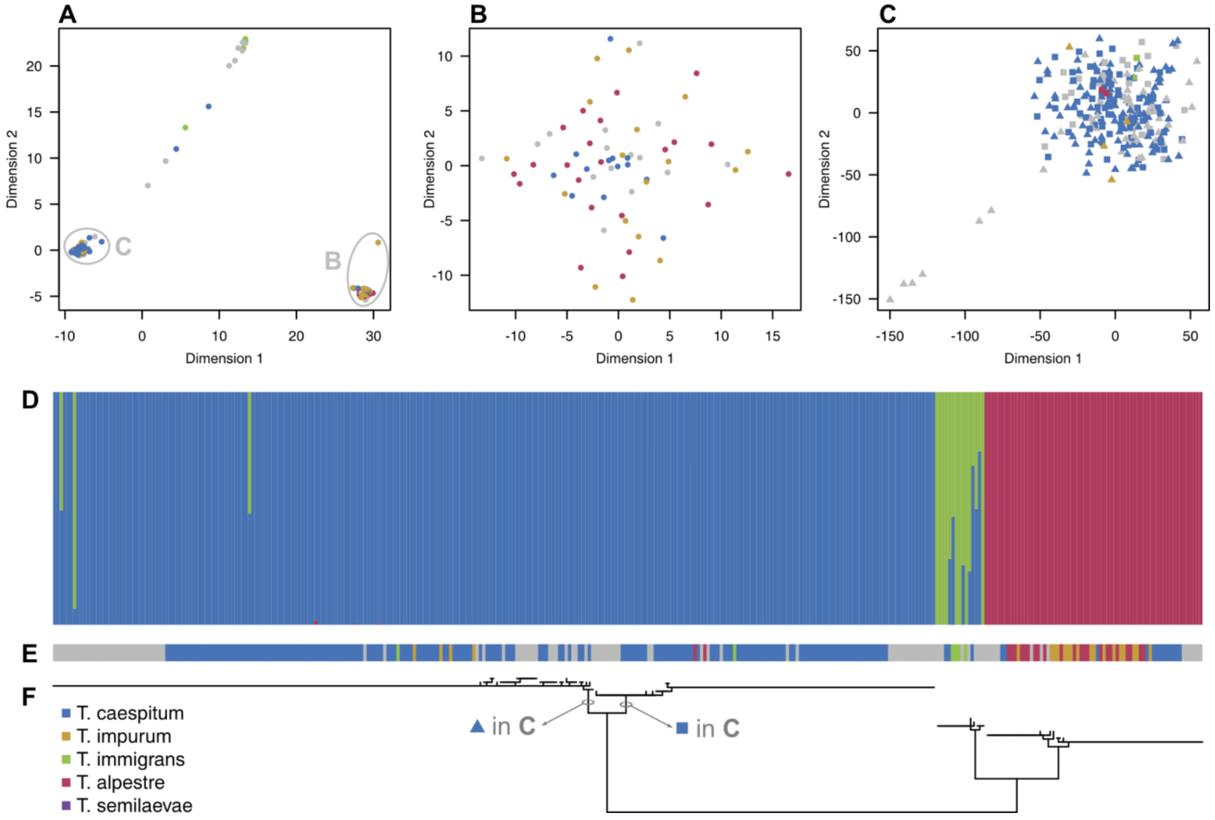


**Figure S7**: Genetic identification of *Tetramorium* samples. **A**: NMDS plot of all samples. The light gray circles indicate clusters of samples reanalyzed separately. Each point is one individual, coloured according to its morphological species identification. Points in grey designate individuals for which morphological identification was unsure. **B**: NMDS plot of the subset of samples of the *T. impurum* / *T. alpestre* group. **C**: NMDS plot of the subset of samples of *T. caespitum*, with point shape according to mitochondrial haplotype (see **F**). **D**: Results of admixture with k = 3. **E**: morphological identification. **F**: phylogeny of the COI gene.

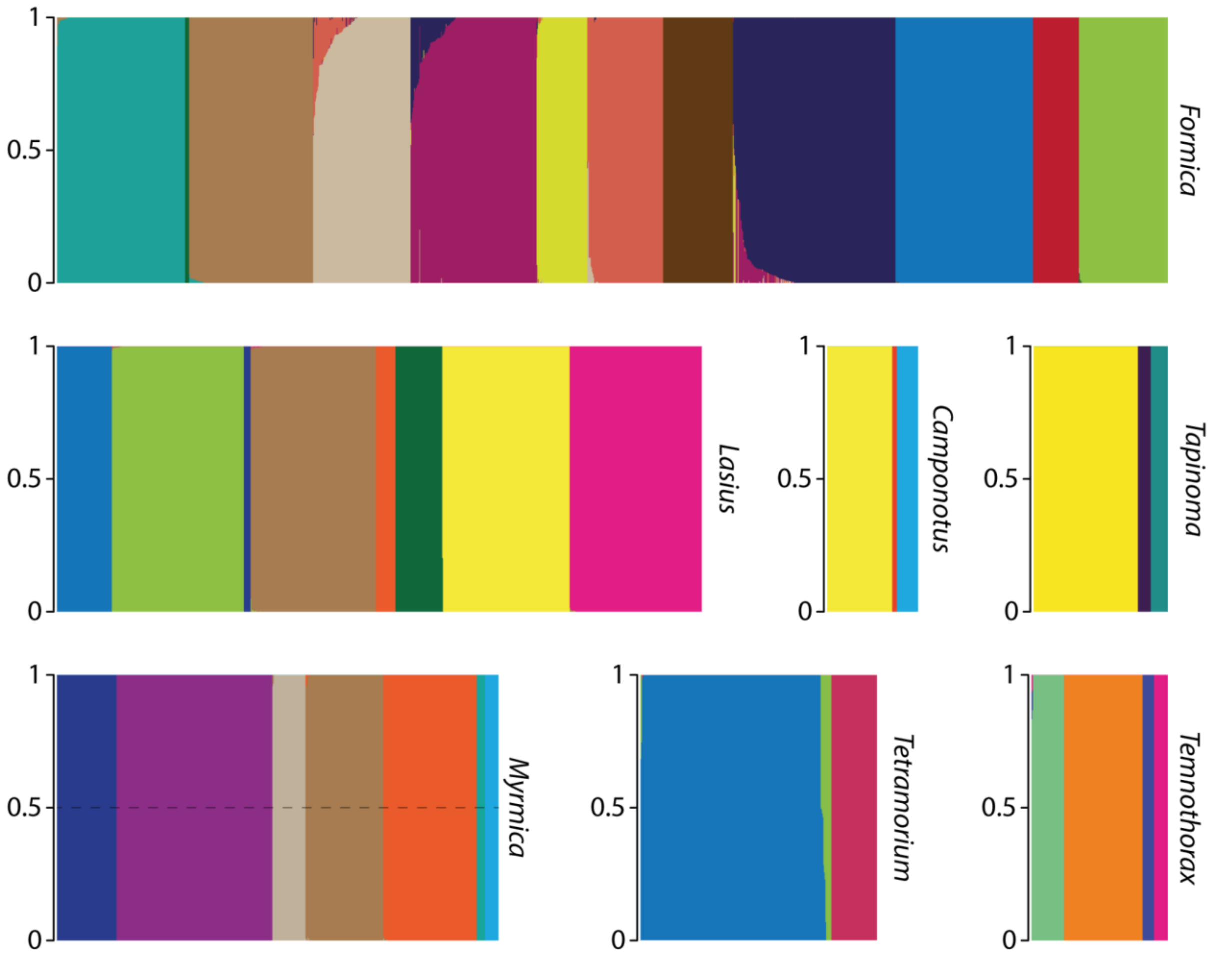


**Figure S8**: Admixture proportion for each of the 4126 retained samples, as in **Figure 1B**, but with species colored as in Figures S1 - S7.

### Analyses of mating phenology and predicted niche

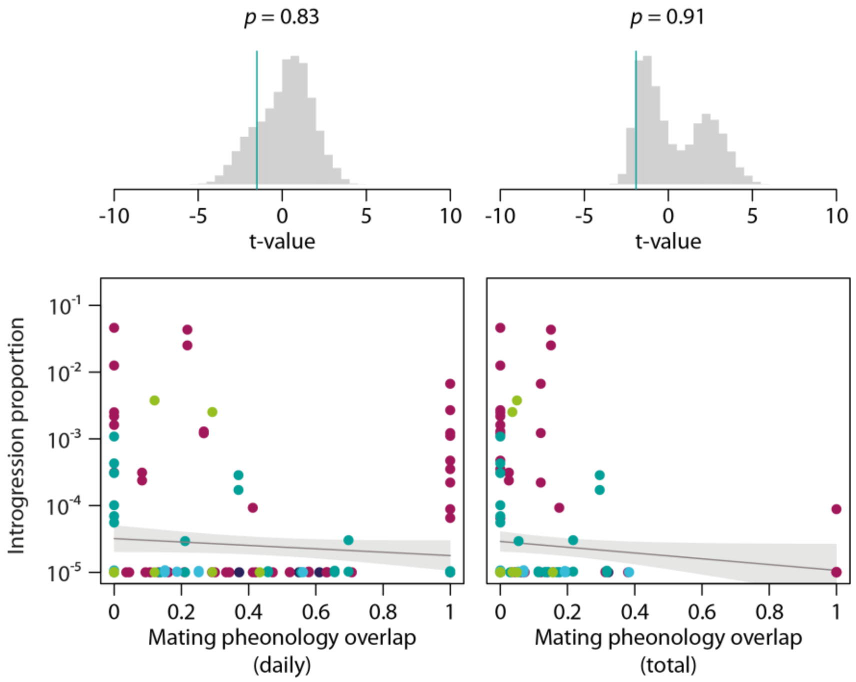

**Figure S9**: Relationship between introgression proportion and overlap mating phenology. **A**: overlap in daily phenology. **B**: overlap in total phenology, estimated as the product of overlap in daily and annual phenology (shown in Figure 3). Colours according to genus, as in **Figure 1**. The predicted values and confidence interval are represented by the grey line and area, respectively. Histograms at the top represent the distribution of simulated *t*-values under permutations. The green line is the observed value. The p-value is the proportion of simulated values falling beyond the observed one (one-tailed tests).

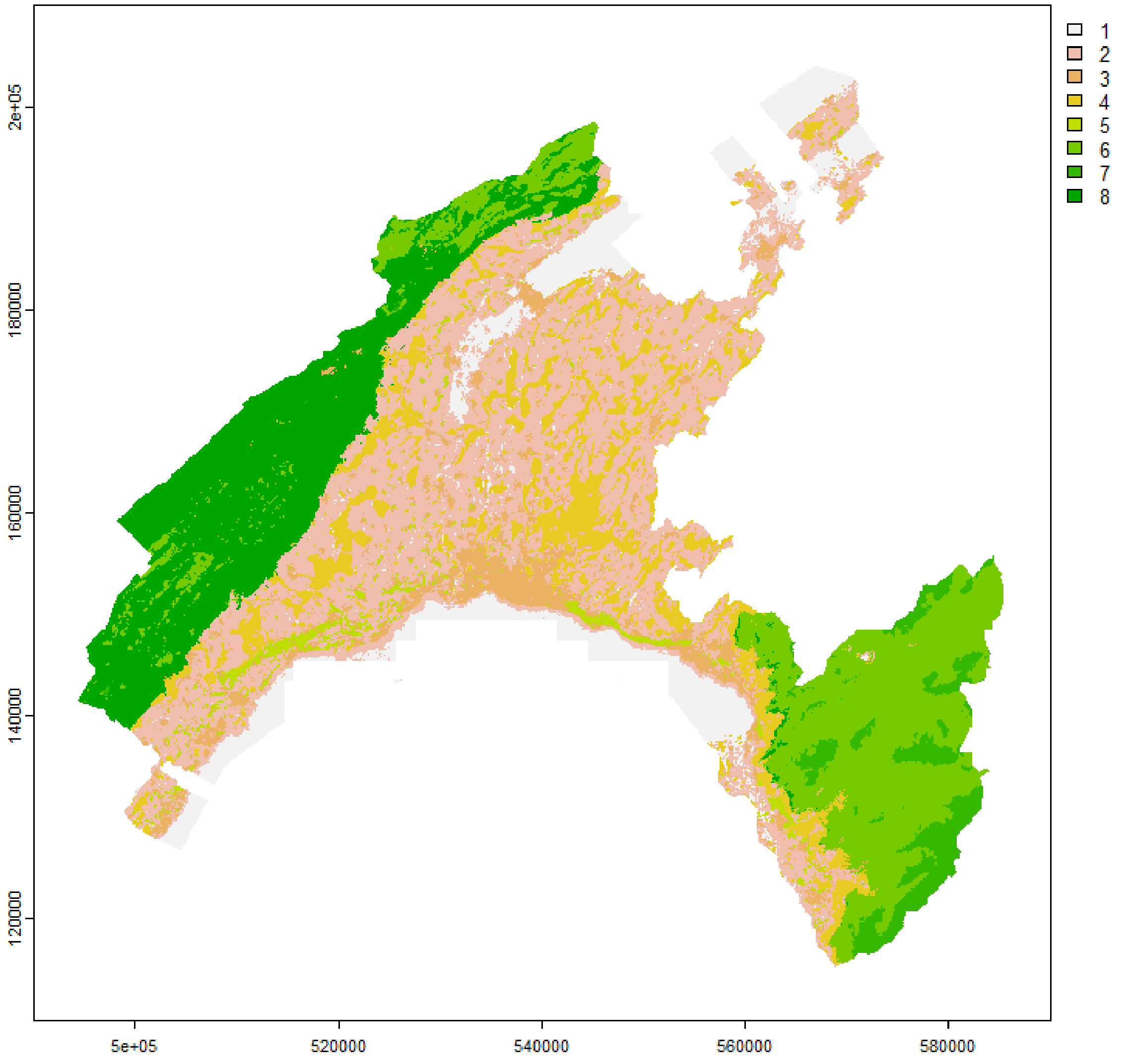


**Figure S10**: map of the study area with the 8 environment types retained for downsampling of the public observations for niche overlap estimations.
